# Preclinical Modeling of Exposure to a Global Marine Bio-Contaminant: Effects of In Utero Domoic Acid Exposure on Neonatal Behavior and Infant Memory

**DOI:** 10.1101/456210

**Authors:** Kimberly S. Grant, Brenda Crouthamel, Caroline Kenney, Noelle McKain, Rebekah Petroff, Sara Shum, Jing Jing, Nina Isoherranen, Thomas M. Burbacher

**Affiliations:** Department of Environmental and Occupational Health Sciences, University of Washington, Seattle, Washington, USA; Center on Human Development and Disability, University of Washington, Seattle, Washington, USA; Washington National Primate Research Center, Seattle, Washington, USA; Department of Pharmaceutics, University of Washington, Seattle, Washington, USA

**Keywords:** domoic acid, prenatal exposure, infant, memory, macaque

## Abstract

Domoic Acid (DA) is a naturally-occurring marine neurotoxin that is increasingly recognized as an important public health issue. Prenatal DA exposure occurs through the maternal consumption of contaminated shellfish/finfish. To better understand the fetal risks associated with DA, we initiated a longitudinal, preclinical study focused on the reproductive and developmental effects of chronic, low-dose oral DA exposure. To this end, 32 adult female Macaca fascicularis monkeys were orally dosed with 0, 0.075 or 0.15 mg/kg/day DA on a daily basis prior to breeding and throughout breeding and pregnancy. The doses included the proposed human Tolerable Daily Intake (TDI) (0.075 mg/kg/day) for DA. Adult females were bred to nonexposed males. To evaluate development during early infancy, offspring were administered a Neonatal Assessment modeled after the human Neonatal Behavior Assessment Scale and a series of Visual Recognition Memory problems using the novelty paradigm. Results indicated that prenatal DA exposure did not impact early survival reflexes or responsivity to the environment. Findings from the recognition memory assessment, given between 1-2 months of age, showed that exposed and control infants demonstrated robust novelty scores when test problems were relatively easy to solve. Performance was not diminished by the introduction of delay periods. However, when more difficult recognition problems were introduced, the looking behavior of the 0.15 mg/kg DA group was random and infants failed to show differential visual attention to novel test stimuli. This finding suggests subtle but significant impairment in recognition memory and demonstrates that chronic fetal exposure to DA may impact developing cognitive processes.

## 1.0 Introduction

Domoic Acid (DA) is a naturally-occurring marine biotoxin that is usually produced by microalgae from the genus *Pseudo-nitzschia*. Mammalian exposures, including human, occur through the consumption of contaminated shellfish and finfish and elevated levels of DA have been documented in ocean waters and coastal regions around the world (Lefebvre and Robertson, 2010, https://www.whoi.edu/science/B/redtide/HABdistribution/HABmap.html.) Based upon an acute poisoning episode involving contaminated mussels on Prince Edward (PEI), Canada in 1987, human symptomology is known to include gastrointestinal distress, confusion, transient and permanent memory loss, coma and death (Perl et al., 1990a, b). Indeed, the clinical label for human DA poisoning, “Amnesic Shellfish Poisoning”, reflects its distinct action on the nervous system that selectively targets cognition. Imaging studies of PEI survivors using Positron Emission Tomography indicated reductions in cerebral glucose metabolism in the hippocampus and the amygdala (Teitelbaum et al, 1990a, b). While the episode of human poisoning in Canada provided data related to the acute effects of DA exposure at very high levels (the estimated exposures ranged from 60 to 290 mg DA or about 1 to 5 mg/kg for a 60 kg person (Perl 1990b, Jefferey et al., 2004), the more subtle effects of sub-toxic, oral DA exposure are largely unknown. Of specific concern are the potential impacts of chronic, low-dose exposure in populations where high shellfish consumption is the dietary norm (Grattan et al., 2016a). Recent results from an environmental epidemiology study focused on chronic, low dose DA exposure reported that the high consumption of razor clams (>15 per week) in adult members of three Native American tribes living on the Western coast of the U.S. was associated with significant memory decrements when compared to a reference group of non-consumers or low-consumers (Grattan et al., 2016b). Deficits in memory were significant enough to impact daily living skills (Grattan et al., 2018).

To date, the developmental neurotoxicity of DA, the focus of this study, has been primarily explored in preclinical rodent models and adverse effects are known to be dose and route dependent (for review see Grant et al., 2010). While the literature base is relatively small, a theme of heightened fetal sensitivity characterized by hippocampal injury and neurological effects can be extracted from the literature (for reviews see Costa et al., 2010; Lefebvre and Robertson, 2010; Doucette et al., 2016). Domoic acid readily crosses the placenta and enters the fetal brain but does not produce structural malformations in exposed pups (Maucher Fuquay et al., 2012). Despite cessation of DA exposure at birth, progressive damage to regions of the hippocampus have been documented over the first 30 days of postnatal life (Dakshinamurti et al., 1993). Results from a number of studies suggest that a single in-utero exposure can cause long-term changes in behavioral development, primarily targeting cognition, social behavior and emotionality and experimental results are frequently accompanied by gender-related differences in performance (Levin et al., 2005, Doucette et al., 2007; Adams et al., 2008; Burt et al., 2008; Tanemura et al., 2009, Ryan et al., 2011; Gill et al., 2012; Marriott et al., 2012, Zuloaga et al., 2016, Shiotani et al., 2017). Early postnatal exposure is associated with a time-dependent profile of neurotoxicity that is characterized by hyperactivity, stereotypic scratching, convulsions and death (Xi et al, 1997). These effects occurred at DA doses 40 times lower than what is required to produce similar clinical symptomology in adult rodents. Reductions in prepulse inhibition (PPI), the normal suppression of the startle response produced by a preceding, non-startling cue (Graham, 1975), have been observed in DA-exposed animals after developmental exposure (Marriot et al, 2012, Adams, 2008). This phenomenon is frequently disrupted in humans with various psychiatric/neurological disorders such as schizophrenia (Takahashi et al., 2011, Haß et al., 2017). To date, there have been no published reports on the effects of prenatal or postnatal DA exposure in human or nonhuman primate infants.

As described above, there is robust evidence from rodent studies that DA can pose significant risks to the fetus but the real-world danger of prenatal exposure in humans has not been explored. Defining these risks to protect human health will depend upon our understanding of this complex chemical when ingested orally at low doses on a chronic basis. This is particularly important because the harmful algal blooms that produce DA are becoming more toxic and widespread as physical processes in ocean waters adapt to warming temperatures (McKibben et al., 2016; Zhu et al., 2017). The research reported in this manuscript is the first to investigate the developmental toxicity of chronic, low-level, oral DA exposure in a nonhuman primate model. The macaque monkey model was selected for this study based on key maternal reproductive and infant developmental similarities shared with humans (Grant and Rice, 2008). The results presented in this paper focus on neonatal behavior and infant visual recognition memory. The dosing levels selected for this investigation were strongly focused on environmental relevance and bracket the proposed human Tolerable Daily Intake (TDI) for DA (Marien, 1996, Costa et al., 2010, Wekell et al., 2004). Results from this investigation provide meaningful translational data on the fetal risks of chronic, low-level maternal DA exposure in a closely-related species.

## 2.0 Materials and Methods

### 2.1 Chemicals and Reagents

DA was purchased from BioVectra (Charlottetown, PE, Canada). Certified calibration solution for DA was purchased from National Research Council Canada (Ottawa, ON, Canada). Optima grade water, methanol, acetonitrile, and formic acid used for bioanalytical assays were purchased from Fisher Scientific (Pittsburgh, PA).

### 2.2 Animal Selection and Study Design

Detailed descriptions of the animals and study design have been reported in Burbacher et al., 2018. Briefly, 32 adult female *Macaca fascicularis* were assigned to either a control group (N=10), a 0.075 mg/kg/day DA group (N=11) or a 0.15 mg/kg/day DA group (N=11) to establish groups with similar average ages and weights. The average age of females was ~ 7 years of age and the average weight was ~3.5 Kg. All females were nulliparous. Three adult males were used for breeding; males were between 7.5 and 8.5 years of age and weighed between 5.4 and 7.9 kg. The study design was based on past studies from our laboratory of in utero exposure to other developmental neurotoxicants such as methylmercury and methanol (Burbacher et al., 1988, 2004) and included a prebreeding, breeding, pregnancy and delivery period. Protocols were implemented to observe signs of DA toxicity (Clinical Observations), detect menstrual bleeding, evaluate general health status (weights, body index) and collect blood for DA analysis without sedation. The adult females had unrestricted access to water and were fed Lab Diet High Protein Monkey Diet biscuits twice a day, once approximately 2 hours before the daily DA dose and once approximately 5 hours after dosing. Animals were housed in individual communicating cages that were equipped with grooming bars that allowed adjacent females contact 24 hours/day. Frozen treats, fresh fruit, music, animated movies and toys were provided to the females on a routine basis. The adult males were housed individually in a separate room away from the females and provided the same suite of enrichments. All animal procedures strictly followed the Animal Welfare Act and the Guide for Care and Use of Laboratory Animals of the National Research Council. Study protocols were approved by the University of Washington Institutional Animal Care and Use Committee to meet the highest standards of ethical conduct and compassionate use of animals in research.

### 2.3 Maternal DA Exposure

Females were dosed daily with DA solution or sweetened water prior to breeding and throughout pregnancy (Jing et al., 2018). Dosing was done in a blinded manner and all dosing solutions were quality controlled and analyzed for DA. To conduct voluntary oral dosing in this study, females were trained with positive reinforcement to drink sweetened water from a syringe offered by the tester. The study protocol required that females continue to be trained until they were reliably drinking from a syringe with no visible spillage. To insure all testers were blind to the dosing history of the animals, solutions containing both DA and 5% sugar water or just 5% sugar water alone were made weekly and stored at 4°C with only animal identifiers on the vials. Each morning, the tester would vortex the vial for approximately 5-10 seconds, and draw 1 ml of solution into a syringe. Dosing occurred at approximately the same time every morning, seven days a week. Dosing was calculated on a mg/kg basis based on weekly weights prior to pregnancy. Males were not exposed to DA.

### 2.4 Offspring

Twenty-eight infants were delivered over a 1 year period, 9 in the control group, 9 in the 0.075 mg/kg/day DA group and 10 in the 0.15 mg/kg/day DA group. Infants were nursery-reared at the Infant Primate Research Laboratory at the University of Washington. This laboratory is equipped to provide 24/7 husbandry and veterinary care for infant monkeys and has set the contemporary standards for the nursery rearing of nonhuman primates (Sackett et al., 2006). Briefly, DA-exposed and control infants were bottle-fed and housed in a temperature-controlled nursery with human incubators until they were able regulate their body temperatures and self-feed. At about 3 weeks of age, animals were transferred to the main rearing quarters and housed in single cages equipped with hanging surrogates and cloth diapers. To facilitate normal social development and future reproductive competence, subjects were placed in stable, mixed-sex groups five days/week in playrooms equipped with a ramp, shelves, swinging chains, toys and a viewing window for tester observations. Several offspring in all of the DA exposure groups were delivered by C-section. Offspring characteristics such as birth weight, crown-rump length and head circumference were similar across the 3 experimental groups (Burbacher et al., 2018). One female in the 0.15 mg/kg/day DA exposure group was vaginally delivered pre-term (<150 days) and had the lowest birthweight at 240 grams. The remaining infants exhibited birthweights and anthropometric measures consistent with normal, full-term newborns for this species.

### 2.5 Maternal and Infant Plasma DA Concentrations

Detailed descriptions related to maternal plasma DA concentrations can be found in Burbacher et al., 2018. Briefly, the plasma DA concentrations 5 hours after dosing were significantly higher for the 0.15 mg/kg/day DA exposure group when compared to the 0.075 mg/kg/day DA exposure group both before and during pregnancy (3.42 ng/ml vs 1.31 ng/ml, before pregnancy; 2.93 ng/ml vs 0.93 ng/ml, during pregnancy). Maternal plasma DA concentrations at delivery were significantly higher than newborn plasma DA concentrations for both DA exposure groups (2.23 ng/ml vs 1.26 ng/ml for the 0.15 mg/kg/day DA exposure group and 1.64 ng/ml vs 0.44 ng/ml for the 0.075 mg/kg/day DA exposure group). Newborn plasma DA concentrations were significantly higher in the 0.15 mg/kg/day DA exposure group compare to the 0.075 mg/kg/day DA exposure group (1.26 ng/ml vs 0.44 ng/ml) (for details, see Burbacher et al., 2018). Plasma DA concentrations decreased after birth and were below the limit of quantification (0.31 ng/ml) by 36 hours of life.

### 2.6 Offspring Assessments

All testers who participated in the evaluation of infants were blind to the exposure history of the animals. Behavioral testers were trained by experienced research personnel and then tested for reliability before collecting data.

#### Neonatal Assessment of Behavior

Our neonatal assessment is adapted from the Neonatal Behavior Assessment Scale (NBAS) used with human infants (Brazelton and Nugent, 2011). The assessment is composed of 5 reflex items and 6 behavioral responses as well as an overall evaluation of the behavioral state of the infant during testing. Tests began with removing the infant from their home-cage and swaddling the infant in a cloth diaper, leaving the upper and lower limbs exposed. Initially, the grasping response of the hands and clasping of the feet were evaluated by placing the index finger against the palm of the hand and sole of the feet. Resistance of the arms and legs to extension was then evaluated by gently but firmly pulling the limbs away from the body. To elicit the rooting and snout reflex, the infant was touched on the corners of the mouth and then on the nose, respectively. Sucking was evaluated by inserting a nipple in the mouth and evaluating the strength of the sucking response. Next, a metal lid was dropped onto a counter to elicit an auditory startle response. The tester then made a lip-smacking noise at the back of each ear to elicit the auditory orienting response. A small colorful toy was held in front of the infant and slowly moved from side to side to examine visual orientation and following. The infant was held above the counter and then quickly moved in the direction of the counter to elicit placing. The arms and legs of the infant were then placed around a rolled-up diaper and strength of grasp evaluated. On the last test item, the infant was placed in a supine position to elicit the righting response. Finally, the tester scored the general behavioral state and temperament of the infant during the evaluation. Offspring were tested every other day during the first 3 weeks of life.

#### Assessment of Object and Social Visual Recognition Memory

A series of recognition memory problems adapted from the Fagan Test of Infant Intelligence (FTII) and the work of Jocelyn Bachevalier at Emory University (Zeamer et al., 2015)were used to evaluate early cognitive processing. Using the paired-comparison paradigm (Fantz, 1956, 1964), this test makes use of the infant’s propensity to prefer novel over familiar visual stimuli and is considered a measure of emerging information processing skills (for review see Burbacher and Grant, 2012). Trials were presented in a preferential looking apparatus where age-graded digitized images were presented on two computer monitors (see Figure 1) at 200 and 210 days of postconceptual (PC) age (roughly equivalent to postnatal ages 40 and 50). Twelve object recognition problems using highly discriminable, colored images of everyday objects were presented at 200 days PC. These test problems also included 1 of 3 delay periods (0, 10 or 30 seconds) between the familiarization period and the test trials. At 210 days PC, 4 social recognition problems using complex images of macaque faces were presented in a second test session.

**Figure 1.**
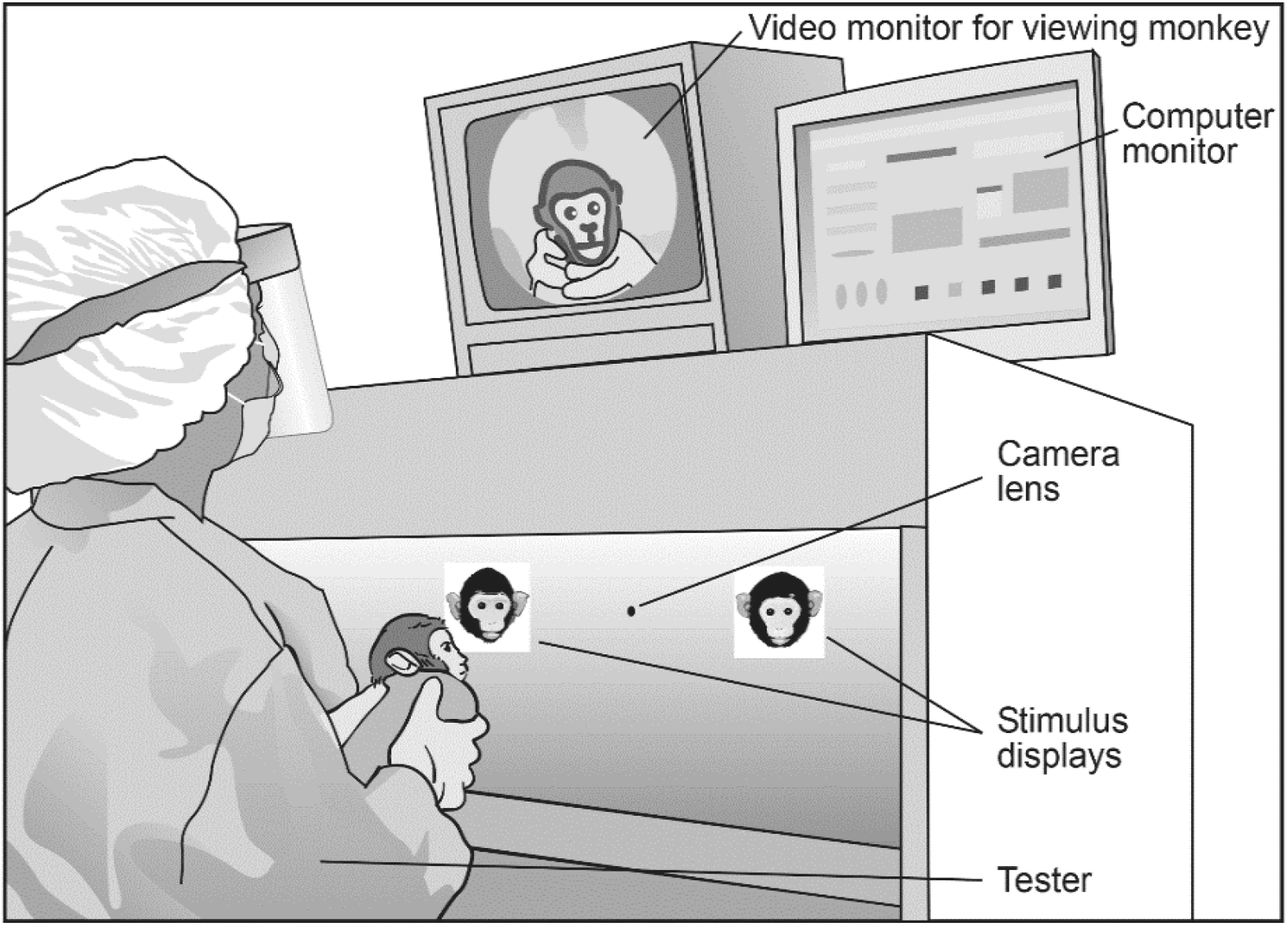
Schematic of Visual Recognition memory test apparatus.

For each trial, the examiner held the swaddled infant in front of the images on the computer monitors and recorded visual fixations to the test stimuli. Looking duration and frequency data were obtained from foot pedals depressed by the tester when the infant actively looked at the right or left stimulus. A physical barrier prevented the examiner from seeing the test stimuli. On each recognition memory problem, the infant was presented with 2 identical visual stimuli until a predetermined amount of looking time had accrued (familiarization time varied from 20 to 30 seconds). When the familiarization period was completed, a 2-part test trial was presented where the previously-viewed stimulus was paired with a novel image. Each part of the test trial (A and B) was 5 seconds in length (10 seconds total) and triggered by the first directed look of the infant to one of the two test stimuli. The position of the familiar and novel stimuli was reversed between Part A and B of the test trial to control for side preferences. At the end of each session, the examiner provided a rating of the subjects overall behavior. Behavior ratings ranged from 1 (asleep or drowsy) to 5 (extremely agitated, untestable). Prior to testing, infants were given a human visual acuity screening test to ensure that each infant had a minimum of 20/800 vision (Teller and Boothe, 1979, Teller, 1997).

### 2.7 Statistical analysis

For the Neonatal Assessment of Behavior test, initial correlational analyses were performed on scores from items that included responses for both hands and feet and right and left limbs. Scores from items with significant correlations (p<0.05) were combined and the average score was included in the analysis for these items. This resulted in scores for 5 reflex items, 6 behavioral-response items and the overall evaluation of attention, irritability and consolability. Scores from these items were combined into an average score for 4 independent factors (Behavioral State, Reflexes, Muscle Tone and Responsivity (see Table 1) in a manner similar to that used in previous studies (Jacobson et al., 1984, Burbacher et al., 1999, Sagiv et al., 2009). Behavioral State was scored from 1 (attentive, cooperative and easy to console) to 4 (inattentive, fussy, and difficult to console). Reflex, Muscle Tone and Responsivity items were scored from weak (1) to strong (4) based on the strength of the response. For the analysis of changes in scores over time, data were averaged over 2-day blocks beginning on day 2 of age. If infants were tested more than once within a 2-day block, the average score over the 2 days was used in the analysis. DA exposure-related effects on factor scores over the first 3 weeks of life were analyzed using a repeated measures analysis of variance (ANOVA), with DA exposure group as the between-subjects factor and age block as the within-subjects factor. To test whether or not sex of the offspring or offspring delivery type (natural or C-section) affected the factor scores over time or interacted with DA exposure to affect scores, two-way repeated measures ANOVAs were used, with DA exposure group and sex (or delivery type) as the between-subjects factors and age block as the within-subjects factor. The results are presented as mean□± 95% confidence intervals. A probability level□<□0.05 was considered significant

**Table 1:** Distribution of Test Items for Neonatal Assessment of Behavior Factors Factor Test Items

| Factors | Factor Test Items |  |  |  |  |
| --- | --- | --- | --- | --- | --- |
| Behavioral state | Attention | Irritability | Consolability |  |  |
| Reflexes | Rooting | Righting | Snout | Placing | Sucking |
| Muscle tone | Resistance | Grasping | Clasping |  |  |
| Responsivity | Startle | Orient<sup>1</sup> | Follow<sup>2</sup> |  |  |

<sup>1</sup> Orient to Auditory and Visual Stimuli
<sup>2</sup> Follow Visual Stimuli

Consistent with the scoring methods used for visual recognition data, the proportion of a subject’s looking time to the novel test stimulus was calculated for each memory problem. For the object recognition problems with delays administered at 190 days PC, novelty scores were calculated for each problem and a mean novelty score was then calculated for the 3 problems within each of the 3 delay periods (0, 10 or 30 seconds). For the social recognition problems given at 200 days PC, novelty scores were calculated for each of the 4 problems and a mean novelty score was then calculated across the problems. The mean novelty scores for each infant are then used to calculate on overall mean score for the DA exposure groups for the different test sessions. While the novelty preference score is the primary outcome of this test, additional variables that provide information regarding patterns of looking/visual attention (e.g. mean duration of visual fixations to targets during familiarization and test periods, mean # gaze shifts between targets divided by exposure time during familiarization and test periods, mean time required time to reach requisite looking time during the familiarization period) can also be extracted from test data. For the current study, data from infants with behavior ratings of 1 (asleep, drowsy) or 5 (untestable) were not used in the analysis.

For the initial analysis, mean novelty scores obtained for each DA exposure group during the first session with the 3 delay period and the second session using social stimuli were used to test the performance of the groups against chance looking (50%), using a single sample t-test. Secondary analyses focused on DA exposure group differences. A repeated measures ANOVA, with DA exposure group as the between-subjects factor and delay period as the within-subjects factor, was used to examine the novelty scores over the 3 delay periods included in the first test session. To test whether or not sex of the offspring or offspring delivery type (natural or C-section) affected the novelty scores over the delays or interacted with DA exposure to affect scores, two-way repeated measures ANOVAs were used, with DA exposure group and sex (or delivery type) as the between-subjects factors and delays as the within-subjects factor. For the second test session using social stimuli, a univariate ANOVA was used to test for DA exposure group differences in novelty scores. Repeated measures and univariate ANOVA models were also performed to examine whether there were differences in the pattern of looking to the stimuli during the familiarization and test periods (see variables listed above). The results for the novelty preference scores are presented as mean□± 95% confidence intervals. A probability level□<□0.05 was considered significant for all tests.

## 3.0 Results

### 3.1 Assessment of Neonatal Behavior

A summary of the results for the 4 factors from the neonatal assessment is shown in Figure 2. Results of the initial repeated measures ANOVA did not indicate a DA exposure effect for any of the 4 factors (p> 0.05, all tests). Scores for 3 of the 4 factors changed with age (Reflexes, Muscle Tone and Responsivity) (p< 0.02, all test). In addition, the results of the two-way repeated measures ANOVAs did not indicate that sex of the offspring or delivery type affected the factor scores directly or that these variables interacted with DA exposure to affect the outcomes (p> 0.05, all tests).

**Figure 2.**
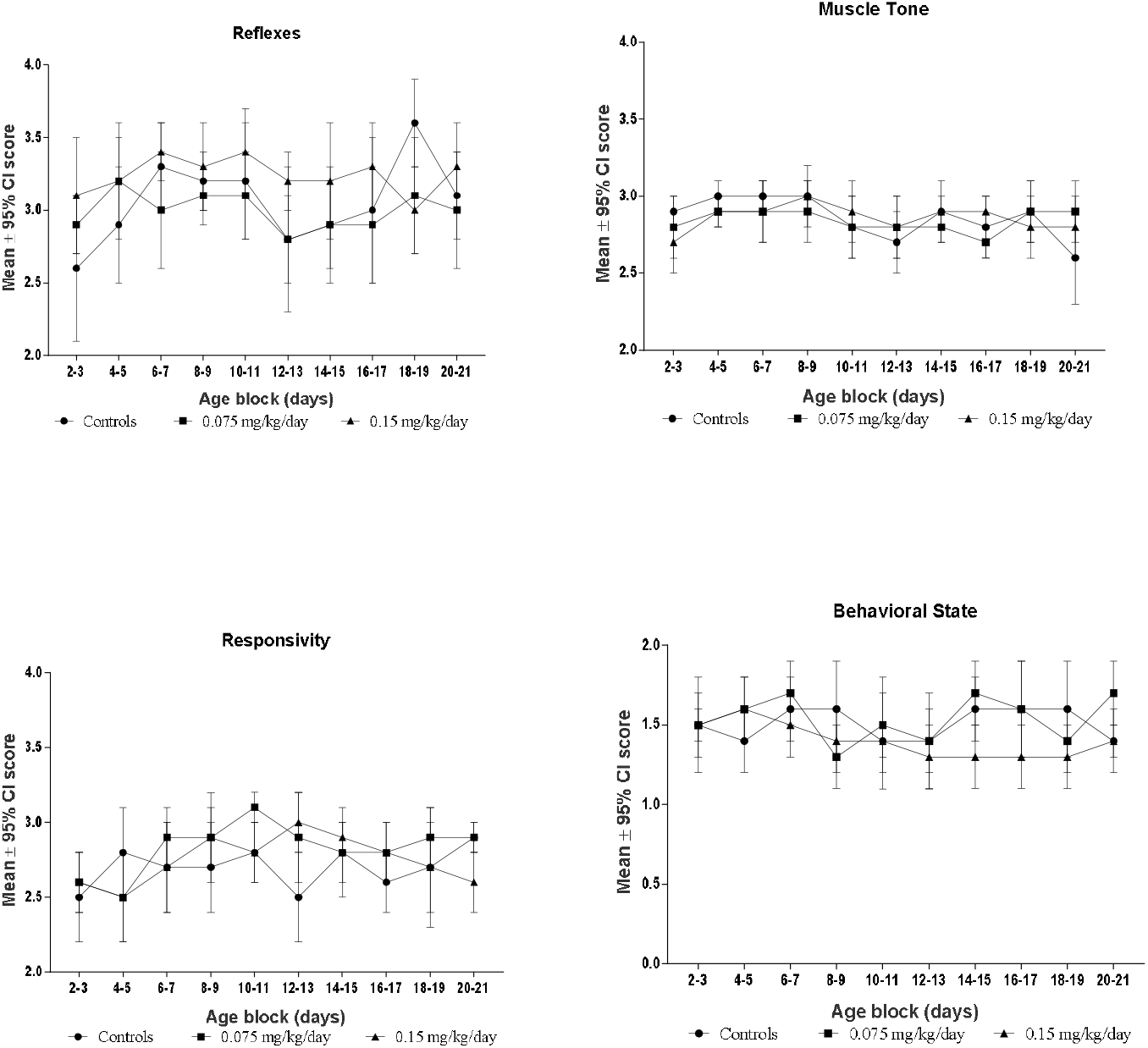
Results of Neonatal Assessment during First 3 Weeks of Postnatal Life

### 3.2 Assessment of Object and Social Recognition Memory

A summary of the novelty scores for the recognition memory assessments is shown in Table 2. For the initial session that included the delay periods, the results of the one-sample t-test indicated that each of the 3 DA exposure groups exhibited mean novelty scores significantly above chance for all of the delays (p<0.05, all tests). For the second session that utilized social recognition problems, the mean novelty scores for the control and 0.075 mg/kg/day DA exposure group on the second test were also significantly above change (p<0.05). However, the mean score for animals in the 0.15 mg/kg/day DA exposure group was not significantly above chance (53.9±3.5 % looking time to novel stimuli, p=0.29, n.s.), indicating random looking behavior and lack of memory for the familiarization stimuli.

**Table 2:** Mean ± SE (Range) Percent Looking Time to Novel Stimuli^1^.

|  | <u>Object Memory Scores</u> |  |  | <u>Social Memory Scores</u> |
| --- | --- | --- | --- | --- |
| DA Exposure Group | 0 Second | 10 Second | 30 Second |  |
| Controls (N=9) | 60.1±4.0<br>(31.7-74.2) | 75.4±3.5<br>(59.2-95.0) | 68.8±5.5<br>(47.8-95.0) | 62.4±4.0<sup>2</sup><br>(44.4-73.3) |
| 0.075 mg/kg/day (N=9) | 60.7±4.4<br>(42.2-85.5) | 66.6±4.3<br>(54.1-85.4) | 70.0±4.9<br>(51.2-96.5) | 63.3±3.5<br>(39.1-72.4) |
| 0.15 mg/kg/day (N=10) | 74.7±4.5<br>(51.4-94.9) | 67.8±4.8<br>(34.0-85.4) | 67.8±6.1<br>(44.0-100.0) | **53.9±3.5\***<br>(34.7-73.3) |

<sup>1</sup> Values are presented as group means ± SE percent time looking at novel stimuli and range of novelty scores.
<sup>2</sup>N=8 (1 infant tested at wrong age, dropped from analysis)
\*No significant novelty preference, $p < .05$

Results from analyses focused on DA exposure group differences did not indicate significant differences in the novelty scores across the 3 DA exposure groups for either test session (object or social) (p>0.05). Sex of the offspring and delivery type also did not affect the novelty scores directly nor did these variables interact with DA exposure to affect the novelty scores for either test (object or social) (p> 0.05, all tests). Finally, results from analyses of the variables focused on pattern of looking during the familiarization and test periods did not indicate significant differences in scores across the 3 DA exposure groups for either test session (object or social) (p>0.05) (data not shown).

## Discussion

This study is the first to investigate the effects of prenatal DA exposure on infant neurodevelopment in a preclinical nonhuman primate model. Neonatal behavioral examinations in the nursery indicated that there were no significant differences between control and DA-exposed infants on Brazelton-derived factors of behavioral state, reflexes, muscle tone and responsivity. Findings from the recognition memory assessment at approximately 35 and 45 days postnatal showed that controls and DA-exposed infants demonstrated significant novelty preferences (memory scores) when test problems were composed of simple, highly-discriminable images of everyday objects. It was only when problems were made more difficult by using complex social stimuli that infants in the 0.15 mg/kg DA group failed to direct a significant proportion of looking time to the novel test stimuli. This finding demonstrates that prenatal DA exposure had a significant effect on recognition memory in the more highly exposed infants. In studies with human infants, problems using social stimuli are considered more difficult (i.e. require greater study time to elicit a comparable novelty response) than problems using images of abstract patterns (Rose, 1980). The sensitivity of social stimuli to individual differences in information processing is widely recognized and the reason that this class of stimuli is employed in human studies with both clinical and research cohorts, including infants exposed to chemicals in-utero.

The novelty test paradigm was selected for use in this study because demonstration of a novelty response requires the ability to discriminate, remember and categorize information. The percent looking time to the novel stimulus during test trials is considered a proxy for recognition memory as some aspects of the familiar stimulus must be encoded in memory for the novelty response to occur. Infant macaque monkeys demonstrate a significant novelty preference by 4-6 weeks of postnatal age and the strength of this response increases over the first several months of life (Gunderson and Sackett, 1984, Bachevalier et al., 1993, Zeamer et al., 2010). Longitudinal work with nonhuman primates has provided insights into the neural substrates of this fundamental memory system (Bachevalier, 2015). While visual recognition memory is dependent on hippocampal functioning, there is mounting evidence that novelty preferences during the first few months after birth are also supported by the perirhinal cortex (e.g. Zeamer & Bachevalier, 2013). The perirhinal cortex is involved in higher cognitive functioning in humans and animals and plays a key role in stimulus familiarity, a foundation of recognition memory Suzuki and Naya, 2014, Brown et al., 2010). A diminished capacity to solve recognition memory problems has been observed in past studies of prenatal neurotoxicant exposure with both human and macaque monkey infants (for review see Burbacher and Grant, 2012). In human infants, reduced novelty preference scores have been associated with prenatal exposure to environmental contaminants such as phthalates (Ipapo et al., 2017), polychlorinated biphenyls (Boucher et al., 2014), methylmercury (Oken et al., 2005), and chlordecone (Dallaire et al., 2012) as well as recreational drugs such as cocaine (Singer et al., 2005, Chiriboga et al., 2007) and alcohol (Jacobson et al., 1985, 2002). Comparative studies with macaque monkeys have demonstrated the sensitivity of this cognitive test paradigm to naturally-occurring perinatal risk factors (Gunderson et al., 1987, 1989), perinatal exposure to environmental tobacco smoke (Golub et al., 2007) and prenatal exposure to neurotoxicants such as methylmercury (Gunderson et al., 1986, 1988), alcohol (Clarren et al., 1992) and methanol (Burbacher et al., 1999).

The underlying basis of the diminished performance observed in the 0.15 mg/kg infants is not known but worth exploring within theoretical and empirical restraints. One possibility is that the infants had difficulty harnessing attention during the familiarization period. If this were the case, their pattern of looking behavior should be different from that seen in the other experimental groups. However, there were no group differences in the mean frequency or duration of visual fixations to the familiarization or test stimuli. There were also no differences between the groups in the time required to reach the pre-determined familiarization criteria. These data suggest that attentional differences are unlikely to be driving the results from this study. Another mechanism that has been proposed to account for individual differences in recognition memory performance is speed of processing. Results from this study are consistent with this theoretical approach. When test images were simple representations of objects, DA-exposed infants demonstrated significant novelty responses even when 10 and 30 second delay periods were introduced. When more complex, less-discriminable test stimuli were used and processing abilities were challenged during the familiarization period, exposed infants were unable to demonstrate recognition. This suggests that the time available during the familiarization period on these problems was not adequate to fully process and encode the test images. Individual differences on human cognitive tests are due to a number of variables but speed of information processing is considered central to performance (Kail, 1994, Fagan, 2000, McFarland, 2017). For example, in a longitudinal birth cohort study conducted in New York, preterm human infants exhibited significantly reduced novelty preferences relative to their full-term counterparts (Rose et al., 2001) but deficits in performance were eliminated when familiarization periods were lengthened to provide more time to process test stimuli (Rose et al., 1980). When children in this cohort were reevaluated on cognitive exams at 11 years of age, structural equation modeling identified speed of processing as one of the attributes of cognition most highly associated with the successful execution of complex mental operations as well as long-term impairments in performance (Rose et al., 2011). Reductions in processing speed have been identified in infants and children after prenatal exposure to neurotoxicants such as alcohol (Burden et al., 2005) and dichlorodiphenyltrichloroethane (DDT) (Gaspar et al., 2015).

Given that DA has demonstrated latent neurotoxicity in other studies, it is reasonable to hypothesize that the effects documented in this study may not fully represent the long-term consequences of prenatal exposure on behavioral development. After early-life chemical or drug exposure, adverse consequences may be transient or attenuated by neural processes that compensate for structural brain damage (Rice and Barone, 2000). Later in life and particularly during the aging process, chemically-induced changes in the brain may be expressed when neural reserves are less able to mask preexisting injuries. Research with humans and multiple animal species has linked DA exposure with the latent onset of neurological effects. This was first identified in a human case of acute DA poisoning where, after a relatively symptom-free year, the subject developed temporal lobe epilepsy and died (Cendes, 1995). At autopsy, this patient showed serious bilateral hippocampal sclerosis. Adult rats exposed to DA also exhibit changing neurological symptomology and after recovery from acute toxicity, enter a period of clinical silence before the onset of progressive, recurrent seizures (Muha and Ramsdell, 2011, Tiedeken and Ramsdell, 2013). In young California sea lions receiving veterinary treatment for DA toxicosis, complete recovery from acute poisoning can be followed by a progressive and frequently lethal epilepsy-like syndrome that is not clinically manifest until months after cessation of exposure (Ramsdell and Gulland, 2014). The authors suggest that epileptic disease is a delayed manifestation of DA poisoning in affected animals. Long-term follow-up of the animals in the present study will provide a unique opportunity to study latent neurotoxicity following prenatal exposure in a highly translational primate model. This is particularly important because general observations of the infants in this cohort provided anecdotal evidence of intentional hand and arm tremors in several DA-exposed infants beginning when they were approximately 2 months of age.

The results described in this manuscript are the first to characterize the effects of prenatal DA exposure on neonatal behavior and visual recognition memory in infant macaque monkeys. While the long-term consequences of early changes in neurodevelopment are not known, it has been cogently argued that in humans, small and even subtle impacts of developmental neurotoxicants, when amortized across populations and generations, can have influential and significant impacts on society (Bellinger, 2012). The animals in this birth cohort are now being evaluated on a suite of developmental exams to study the effects of DA on learning and memory, social behavior, temperament and EEG activity during late infancy and pre-adolescence. Results from these assessments will be critical to defining the neurodevelopmental risks posed by fetal DA exposure at levels close to the human TDI and will identify basic domains of behavior and health that are vulnerable to long-term disruption.

## Acknowledgments

This research was supported by grants from the U.S. National Institutes of Health R01 ES023043, P51 OD010425, HD083091 and NCATS Grant TL1 TR000422 (SS). The authors would like to acknowledge the dedicated staff and volunteers of the Infant Primate Research Laboratory and the University of Washington National Primate Research Center for their assistance with the care and support of this cohort. In addition, we would like to thank Mr. Steven Ellis for his assistance with data processing and analysis.

